## Supplementary Material for "New insights into large tropical tree mass and structure from direct harvest and terrestrial lidar"

This document contains two appendices to our paper entitled 'New insights into large tropical tree mass and structure from direct harvest and terrestrial lidar'. Appendix A considers the allometric approach to estimating the above-ground biomass of the four harvested trees. In particular, we consider our choice of allometric model. Appendix B then considers the lidar-derived approach to estimating above-ground biomass. We demonstrate how the various data processing steps influence the performance of estimates.

**Appendix A: Allometric model selection**

The two-parameter pan-tropical model described in Chave et al. [1] (equation 4, predictor variables: stem diameter, $D$ (cm), tree height, $H$ (m), and basic wood density, $\rho$ ($\mathrm{g\,cm^{-3}}$)) was used in the paper to represent the allometric approach. Our justification for selecting this particular model is that it provided some of most accurate predictions of above-ground biomass for the four harvested trees, relative to other regional and pan-tropical models, and because of its wide application throughout the literature. Here, we demonstrate this accuracy by generating allometric estimates of above-ground biomass, $\mathrm{AGB_{est}}$, using the four alternative models described below, and compare these estimates with the harvest-derived reference values, $\mathrm{AGB_{ref}}$. It is noted that in most instances, the following models were published in their respective papers in log-space, but for consistency, are presented here in real-space. For the avoidance of doubt, here, each incorporates a correction factor (based on the standard error of the regression, $\sigma$), to compensate for bias in $\mathrm{AGB_{est}}$ arising from differences in the expectations of the log-normal and normal distributions [2].

(1) The regional model described in Chambers et al. [3], constructed using data collected from 315 harvested tropical trees located approximately 60 km north of Manaus, Brazil. This model uses the single predictor variable, $D$, and takes the form:

$$\mathrm{AGB_{est}} = e(\frac{0.297^2}{2})\ e(-0.370 + 0.333\ln(D) + 0.933[\ln(D)]^2 - 0.122[\ln(D)]^3)$$

(2) The first of two pan-tropical models described in Feldpausch et al. [4], constructed using data collected from 1816 tropical trees harvested from across the tropics. This model shares some data with those used in the Chave et al. [1] model. Considered are the predictor variables $D$ and $\rho$, and the model takes the form:

$$\mathrm{AGB_{est}} = e(\frac{0.360^2}{2})\ e(-1.822 + 2.334\ln(D) + 0.163[\ln(D)]^2 - 0.025[\ln(D)]^3 + 0.979\ln(\rho))$$

(3) The second pan-tropical model described in Feldpausch et al. [4], constructed using the same data. This model further includes the predictor variable $H$, and takes the form:

$$\mathrm{AGB_{est}} = e(\frac{0.322^2}{2})\ e(-2.921 + 0.989\ln(D^2H\rho))$$

(4) The alternative pan-tropical model described in Chave et al. [1, 5], constructed using data collected from 4004 tropical trees harvested from across the tropics. This model considers

the predictor variables $D$, $\rho$ and $E$, and takes the form:

$$\text{AGB}_{\text{est}} = e(-2.024 - 0.896E + 0.920\ln(\rho) + 2.795\ln(D) - 0.046[\ln(D)]^2) \tag{1}$$

Where $E$ is a measure of environmental stress, and at the location of our site, $E = -0.054$.

Table 1 below replicates upper table 2 in the paper, comparing $\text{AGB}_{\text{est}}$ predicted from these models, with $\text{AGB}_{\text{ref}}$, and also reports on tree-scale error and up-scaled error (i.e., the cumulative AGB of the four trees) in these estimates. It is observed the only model providing estimates with accuracy similar to the selected model is (3).

| **ID** | **AGB$_{ref}$** (kg) | **(1) Chambers et al. (2001)** $[D]$ | | | **(2) Feldpausch et al. (2012)** $[D, \rho]$ | | | **(3) Feldpausch et al. (2012)** $[D, H, \rho]$ | | | **(4) Chave et al. (2014)** $[D, \rho, E]$ | | |
|---|---|---|---|---|---|---|---|---|---|---|---|---|---|
| | | **AGB$_{est}$** (kg) | **Error** (kg) | **R. E.** (%) | **AGB$_{est}$** (kg) | **Error** (kg) | **R. E.** (%) | **AGB$_{est}$** (kg) | **Error** (kg) | **R. E.** (%) | **AGB$_{est}$** (kg) | **Error** (kg) | **R. E.** (%) |
| T1 | 3960.1 | 4627.2 | 667.1 | 16.8 | 4779.8 | 819.7 | 20.7 | 3571.4 | −388.7 | 9.8 | 4255.1 | 295.0 | 7.5 |
| T2 | 18 584.2 | 10 404.6 | −8179.6 | 44.0 | 25 691.6 | 7107.4 | 38.2 | 24 399.1 | 5814.9 | 31.3 | 23 507.0 | 4922.8 | 26.5 |
| T3 | 8392.6 | 7751.5 | −641.1 | 7.6 | 11 820.4 | 3427.8 | 40.8 | 9120.9 | 728.3 | 8.7 | 10 600.4 | 2207.8 | 26.3 |
| T4 | 5521.1 | 5253.5 | −267.6 | 4.8 | 5706.0 | 184.9 | 3.3 | 4874.4 | −646.7 | 11.7 | 5085.2 | −435.9 | 7.9 |
| Mean | - | - | - | 18.3 | - | - | 25.8 | - | - | 15.4 | - | - | 17.0 |
| Summed | 36 458.0 | 28 036.8 | −8421.2 | 23.1 | 47 997.8 | 11 539.8 | 31.7 | 41 965.8 | 5507.8 | 15.1 | 43 447.7 | 6989.7 | 19.2 |

Table 1: Performance of estimates of above-ground biomass (AGB$_{est}$) from four regional and pan-tropical allometric models. Error in these estimates is calculated from comparison with directly measured reference values (AGB$_{ref}$). R. E. denotes relative error. Mean relative error is the average of tree-scale relative errors. Summed values refer to the cumulative AGB of the four trees. For those models using basic wood density as a predictor variable, we used the mass-weighted estimate of above-ground basic woody tissue density, which is defined in the methods section of the paper.

**Appendix B: Processing terrestrial lidar data**

Lidar-derived estimates of tree-scale above-ground biomass, $AGB_{est}$, were calculated from estimates of above-ground green woody volume and above-ground basic woody tissue density. Retrieving these estimates of volume from the raw lidar data required the following sequential steps:

(1) Individual scans were registered onto a common coordinate system using *RiSCAN PRO* (v2.7.0) [6].

(2) Point clouds representing the individual trees were extracted from the larger-area point clouds using *treeseg* (v0.2.0) [7].

(3) Points were classified as returns from either wood or leaf surfaces using *TLSeparation* (v1.2.1.5) [8].

(4) Points from buttresses were manually removed using *CloudCompare* (v.2.10.3) [9].

(5) Quantitative structural models (QSMs) representing each tree were constructed from the point clouds using *TreeQSM* (v2.3.2) [10]. The input parameters to *TreeQSM* were automatically selected using *optqsm* (v0.1.0).

Here, we demonstrate the importance of steps (3) and (4) to the performance of $AGB_{est}$. This is undertaken by calculating $AGB_{est}$ from QSMs constructed using: i) the full point clouds after extraction and downsampling via *treeseg*, ii) the so-called 'leaf-off' point clouds after classification via *TLSeparation*, and iii) these leaf-off point clouds after the removal of points from buttresses using *CloudCompare*. These series of tree-level point clouds and corresponding QSMs are illustrated in figures 1 and 2 respectively. Table 2 then compares $AGB_{est}$, retrieved from these sets of QSMs, with the harvest-derived reference values, $AGB_{ref}$, and also reports on tree-scale error and up-scaled error (i.e., the cumulative AGB of the four trees) in these estimates. It is observed that the mean tree-scale relative error in $AGB_{est}$ derived from i), ii) and iii) was 42 %, 5 % and 3 % respectively. That is, for these particular data, the removal of leaf returns to prevent their interference in woody reconstruction was critical to the accuracy of $AGB_{est}$.

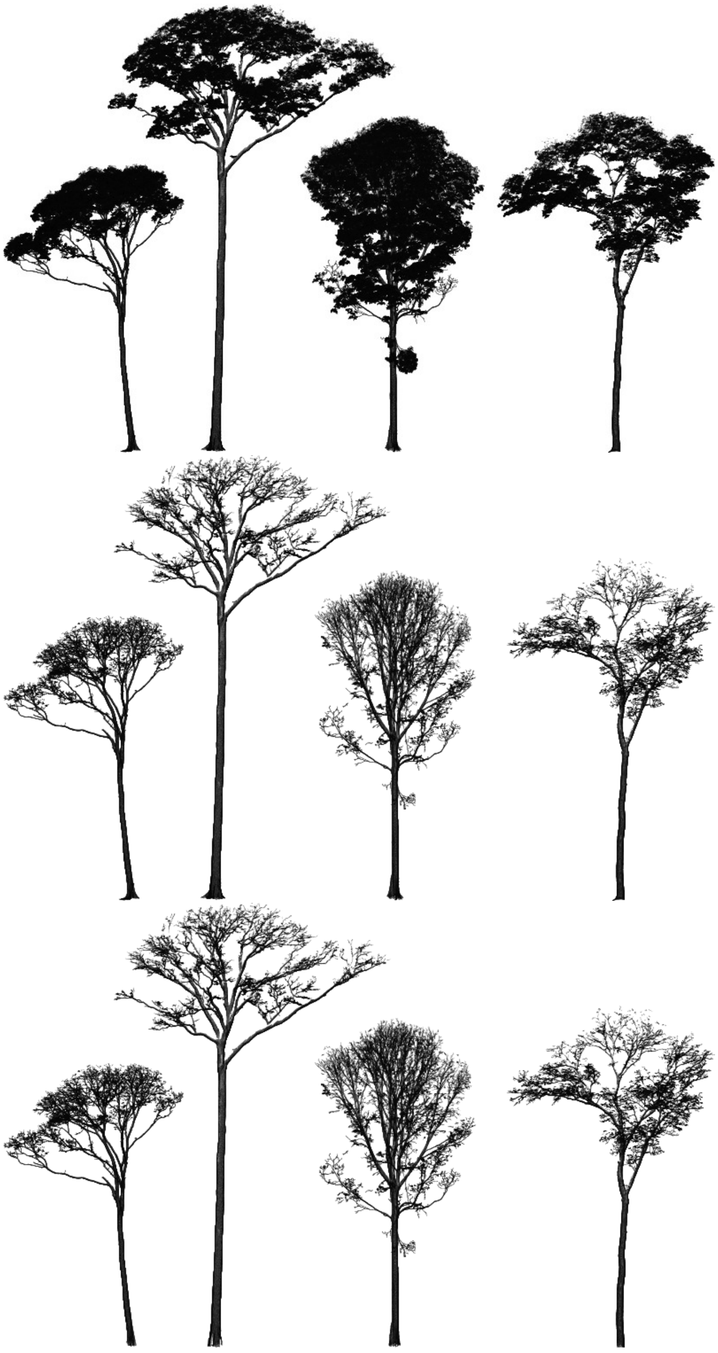

Figure 1: Point clouds of the four harvested trees. To scale; left to right: T1-T4. The top set of clouds were extracted from the larger-area point clouds using *treeseg*. The leaf-off clouds shown in the middle set were generated using *TLSeparation*. The bottom set of clouds show these leaf-off point clouds after points from buttresses were manually removed using *CloudCompare*.

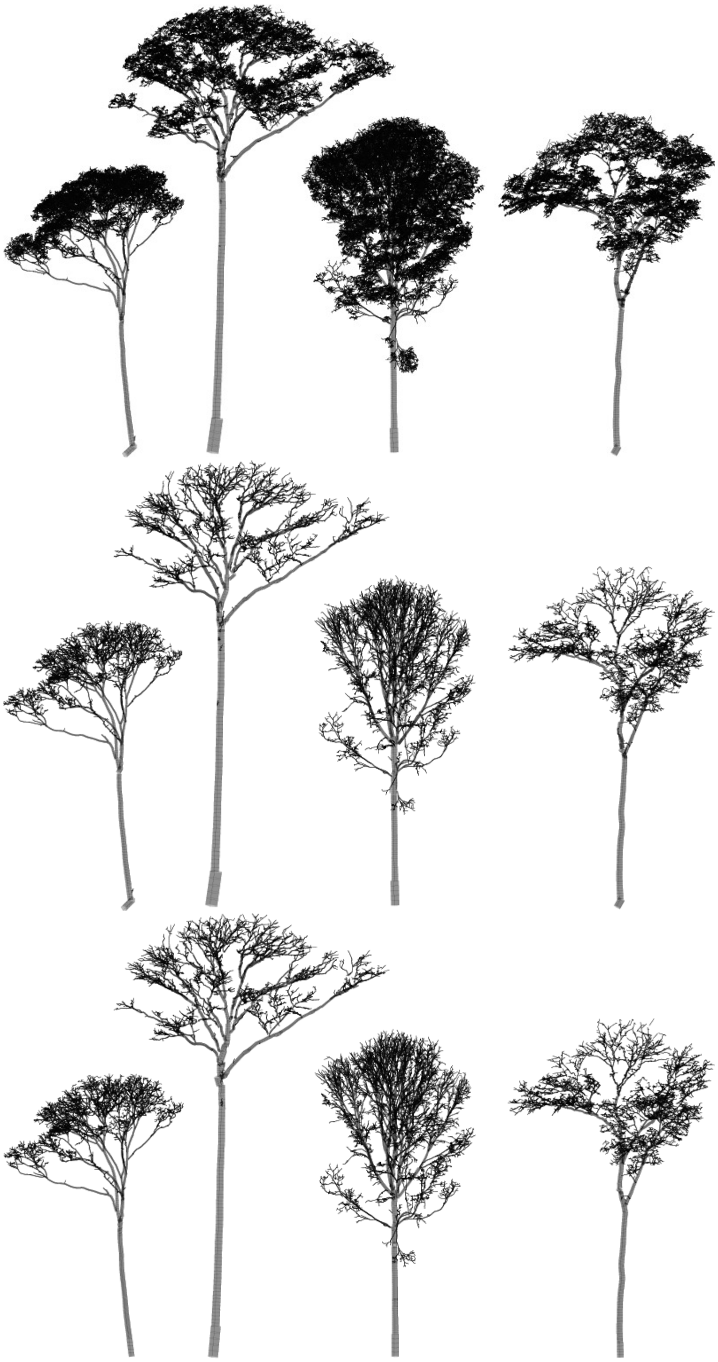

Figure 2: Quantitative structural models constructed from the corresponding point clouds shown in figure 1. To scale, left to right: T1-T4.

| ID | $AGB_{ref}$ (kg) | **(i) Full point clouds** | | | **(ii) Leaf-off point clouds** | | | **(iii) Leaf-off & buttress removal** | | |
|---|---|---|---|---|---|---|---|---|---|---|
| | | $AGB_{est}$ (kg) | **Error** (kg) | **R. E.** (%) | $AGB_{est}$ (kg) | **Error** (kg) | **R. E.** (%) | $AGB_{est}$ (kg) | **Error** (kg) | **R. E.** (%) |
| T1 | 3960.1 | 5516.8 | 1556.7 | 39.3 | 4186.5 | 226.4 | 5.7 | 3673.8 | −286.3 | 7.2 |
| T2 | 18 584.2 | 24 853.1 | 6268.9 | 33.7 | 19 878.7 | 1294.5 | 7.0 | 18 414.5 | −169.7 | 0.9 |
| T3 | 8392.6 | 13 604.4 | 5211.8 | 62.1 | 8957.7 | 565.1 | 6.7 | 8604.6 | 212.0 | 2.5 |
| T4 | 5521.1 | 7399.8 | 1878.7 | 34.0 | 5471.4 | −49.7 | 0.9 | 5485.1 | −36.0 | 0.7 |
| Mean | - | - | - | 42.3 | - | - | 5.1 | - | - | 2.8 |
| Summed | 36 458.0 | 51 374.1 | 14 916.1 | 40.9 | 38 494.3 | 2036.3 | 5.6 | 36 177.9 | −280.1 | 0.8 |

Table 2: Performance of estimates of above-ground biomass ($AGB_{est}$) derived from the QSMs shown in figure 2, and the mass-weighted estimates of above-ground basic woody tissue density (this variable is defined in the methods section of the paper). Error in these estimates is calculated from comparison with directly measured reference values ($AGB_{ref}$). R. E. denotes relative error. Mean relative error is the average of tree-scale relative errors. Summed values refer to the cumulative AGB of the four trees.
